# The effect of vascular health factors on white matter microstructure mediates age-related differences in executive function performance

**DOI:** 10.1101/2020.05.14.096677

**Authors:** David A. Hoagey, Linh T.T. Lazarus, Karen M. Rodrigue, Kristen M. Kennedy

## Abstract

Even within healthy aging, vascular risk factors can detrimentally influence cognition, with executive functions (EF) particularly vulnerable. Fronto-parietal white matter (WM) connectivity in part, supports EF and may be particularly sensitive to vascular risk. Here, we utilized structural equation modeling in 184 healthy adults (aged 20-94 years of age) to test the hypotheses that: 1) fronto-parietal WM microstructure mediates age effects on EF; 2) higher blood pressure (BP) and white matter hyperintensity (WMH) burden influences this association. All participants underwent comprehensive cognitive and neuropsychological testing including tests of processing speed, executive function (with a focus on tasks that require switching and inhibition) and completed an MRI scanning session that included FLAIR imaging for semi-automated quantification of white matter hyperintensity burden and diffusion-weighted imaging for tractography. Structural equation models were specified with age (as a continuous variable) and blood pressure predicting within-tract WMH burden and fractional anisotropy predicting executive function and processing speed. Results indicated that fronto-parietal white matter of the genu of the corpus collosum, superior longitudinal fasciculus, and the inferior frontal occipital fasciculus (but not cortico-spinal tract) mediated the association between age and EF. Additionally, increased systolic blood pressure and white matter hyperintensity burden within these white matter tracts contribute to worsening white matter health and are important factors underlying age-brain-behavior associations. These findings suggest that aging brings about increases in both BP and WMH burden, which may be involved in the degradation of white matter connectivity and in turn, negatively impact executive functions as we age.

## 1. Introduction

One major research focus in investigations of age-related declines in cognitive performance is in the domain of executive function (EF; Goh, An, & Resnick, 2012). EF, an umbrella term encompassing a wide range of higher-order cognitive abilities associated with planning, flexibility/switching, updating, and inhibition that are engaged during complex and novel tasks (Goldstein, Naglieri, Princiotta, & Otero, 2014; McKenna, Rushe, & Woodcock, 2017; Shallice, 1988; Stuss, 1992), declines precipitously with advanced age (Lustig & Jantz, 2015; Spreng, Shoemaker, & Turner, 2017). Given the abundance of cognitive processes attributed to EF, various models have been developed to identify and isolate unique aspects of EF to better understand performance differences observed in aging. Component analyses and structural modeling are often employed to create separation and specificity among cognitive tasks measuring a broad range of EF constructs. However, the reliability and replicability of this work across studies has been questioned, particularly regarding the influence of processing speed (PS), and how delays in reaction or response time measures are entangled with measures of EF (Salthouse, 1991, 1993, 2005; Salthouse & Madden, 2013), and are difficult to disentangle from within a single cognitive assessment test (Salthouse, 2011). Variance in speeded tasks is often shared with aspects of flexibility, shifting, inhibition, decision making, and updating; factor models have been an effective way of teasing apart these confounding measures (Bettcher et al., 2016; Genova, DeLuca, Chiaravalloti, & Wylie, 2013; Henninger, Madden, & Huettel, 2010).

Despite these challenges, research has consistently highlighted the importance of brain health in maintaining high levels of EF performance. For years, research into the effects of lesions and damage to the frontal lobes have pointed to the significance of the prefrontal cortex in EF performance (Alvarez & Emory, 2006; Goldberg, 2002; Jurado & Rosselli, 2007; Lezak, Howieson, Loring, & Fischer, 2004; Miller & Cohen, 2001; Stuss & Alexander, 2000; Stuss & Benson, 1984). However, as pointed out by Stuss (2011), the frontal lobes are large and composed of a diverse set of architectonically unique regions, all of which demonstrate structural and functional connectivity with one another and across the entire cortex (Stuss, 2011). With the advent of more advanced neuroimaging techniques, allowing for *in vivo* exploration of the inter-connections within the prefrontal and among the more distal parietal and subcortical brain regions, the mechanisms responsible for successful EF processing can be more clearly elucidated.

A considerable body of functional neuroimaging research illustrates that while frontal lobe regions activate during specific EF tasks, a larger network of parietal regions are recruited depending on task demands (Buchsbaum, Greer, Chang, & Berman, 2005; Collette & Van der Linden, 2002; Kim, Cilles, Johnson, & Gold, 2012; Owen, McMillan, Laird, & Bullmore, 2005; Ravizza & Carter, 2008; Simmonds, Pekar, & Mostofsky, 2008; Swick, Ashley, & Turken, 2011). Additionally, temporal, occipital, and subcortical regions can play a role in EF, particularly in tasks that rely heavily on working memory, language, motor, or visual processes (Jurado & Rosselli, 2007; Kennedy & Raz, 2009a; Lewis, Dove, Robbins, Barker, & Owen, 2004; Monchi, Petrides, Strafella, Worsley, & Doyon, 2006; Swick et al., 2011). Using resting state functional measures, network-like properties have been identified across many of these same regions found in task-based neuroimaging studies of EF (Cole et al., 2013; Damoiseaux et al., 2006; Madden et al., 2017; Niendam et al., 2012; Nyhus & Barceló, 2009; Reineberg & Banich, 2016; Shirer, Ryali, Rykhlevskaia, Menon, & Greicius, 2012; S. Zhang & Li, 2012). Structural correlates of EF performance have also been reported, most notably, in the frontal and parietal regions of the brain, corroborating task-based functional patterns of activation (Burzynska et al., 2012; H. R. Smolker, Friedman, Hewitt, & Banich, 2018; Weise, Bachmann, Schroeter, & Saur, 2019; Yuan & Raz, 2014).

Given the broad range of functional and morphologic associations, as well as the network-like organization, particularly between frontal and parietal regions, efficient EF performance is hypothesized to be dependent upon the health of connections within and between the frontal and parietal white matter. In fact, there is evidence that white matter volume and diffusion measures represent the majority of the age-related variance attributed to EF performance when combined as part of a multi-modal component analysis, leading Fjell and coauthors to the conclusion that “the major part of the age-related reductions in executive function can be attributed to micro- and macrostructural alterations in brain connectivity” (Fjell, Sneve, Grydeland, Storsve, & Walhovd, 2016). Additional support for the role of white matter-derived associations with EF are found throughout the diffusion imaging literature, emphasizing the importance of white matter connecting frontal and parietal regions via the superior longitudinal fasciculus (SLF; Gallen, Turner, Adnan, & D’Esposito, 2016; Sasson, Doniger, Pasternak, Tarrasch, & Assaf, 2013; H. Smolker, Depue, Reineberg, Orr, & Banich, 2015; H. R. Smolker et al., 2018; J. Zhang et al., 2019), the corpus callosum (Bettcher et al., 2016; Kennedy & Raz, 2009a; Voineskos et al., 2012; J. Zhang et al., 2019), inferior frontal occipital fasciculus (IFOF; H. Smolker et al., 2015), uncinate fasciculus (UF), inferior longitudinal fasciculus (ILF), fornix (Sasson et al., 2013), and various other pathways defined using functional or structural ROIs (Charlton et al., 2006; Grieve, Williams, Paul, Clark, & Gordon, 2007; Kennedy & Raz, 2009a; Shen et al., 2019). However, given the inherent complexity in tasks measuring EF, white matter pertaining to the fornix, UF, posterior corpus callosum, and ILF contribute the most when memory, motor, and language-based tasks of EF are employed (Kennedy & Raz, 2009a; Sasson et al., 2013; J. Zhang et al., 2019). Additionally, reduction in PS is also linked to white matter health in many of these same regions, and covaries with EF performance (Bucur et al., 2008; Deary et al., 2006; MacPherson et al., 2017; Madden et al., 2004; Sullivan, Adalsteinsson, & Pfefferbaum, 2005; Tuch et al., 2005), highlighting the importance of accounting for PS in the study of white matter and EF (Genova et al., 2013).

There is overwhelming evidence that white matter health underlies cognitive performance differences across the lifespan. However, vascular components, such as hypertension and cerebral small vessel disease, evidenced by leukoaraiosis or white matter hyperintensities (WMH), contribute significant variance in explaining the extent and longitudinal progression of these brain-behavior associations (Debette & Markus, 2010), particularly in aging (Raz, Rodrigue, Kennedy, & Acker, 2007). Not only do overall increases in WMH contribute to poorer overall cognitive and EF ability (Au et al., 2006; de Groot et al., 2001; Gunning-Dixon & Raz, 2000; Kloppenborg, Nederkoorn, Geerlings, & van den Berg, 2014; Meier et al., 2014; Nordahl et al., 2006; Rizvi et al., 2020), but the association is regionally specific where WMH location is predictive of specific declines in EF, over and above total WMH burden (Biesbroek, Weaver, & Biessels, 2017; Lampe et al., 2019; Smith et al., 2011). Moreover, some researchers propose that detrimental effects on cognition are specifically due to the vascular nature of WMH, in particular differences in underlying blood pressure, or a more direct influence of increased blood pressure on the health of white matter tracts, as exhibited through associations with white matter diffusion properties (Anstey & Christensen, 2000; Kennedy & Raz, 2009b; Madden, Bennett, & Song, 2009; Maillard et al., 2013; Rizvi et al., 2020; Smith et al., 2011; Vernooij et al., 2008; Waldstein, Giggey, Thayer, & Zonderman, 2005). Thus, it is essential to quantify regional white matter hyperintensities and include blood pressure measurements in the study of white matter and cognitive aging.

Much of the current literature attempts to assess the significance of each of the aforementioned factors using separate, univariate regressions. This approach is restrictive in that it fails to account for the interconnectedness of the brain and the myriad of health factors that may influence cognitive outcomes. Given the complex interplay of both brain and health factors that influence cognition in aging, a multitude of variables must be accounted for and modeled with appropriate specificity to elucidate underlying mechanisms. A multivariate statistical modeling technique such as structural equation modeling, can be used to demonstrate how a “disconnection” can emerge if estimates of brain health are shown to mediate and account for the variance in cognitive differences in aging. As discussed above, differences in white matter diffusion likely reflect altered communication efficiency between the frontal and parietal interconnecting fiber pathways, and could thus lead to a disconnection of cortical communication and poorer cognitive performance in aging adults. Disconnection models have previously shown that efficient communication among higher-order cognitive centers is required for optimal cognitive processing with aging (Antonenko & Flöel, 2014; Bartzokis et al., 2004; Bennett & Madden, 2014; Fjell et al., 2016; Madden et al., 2017; O’Sullivan et al., 2001). To test for white matter tract type specificity, the projection fibers comprising the corticospinal tract is contrasted with frontal white matter association tracts. Projection fibers develop early and are last to show effects of aging, whereas frontal and posterior parietal white matter have the most protracted development and earliest aging vulnerability. This notion has been referred to as a retrogenesis of white matter regions during the course of aging (Raz, 2000; Salat et al., 2004). Combined, this statistical and conceptual framework of modeling disconnection allows for variance among a large set of variables to be attributed to specific pathways, leading to a more refined conceptualization of the neural mechanisms involved in cognitive aging than what is currently understood.

The current study sought to investigate these ideas in an aging sample by modeling variable estimates of EF, PS, white matter diffusion, white matter hyperintensity burden, and blood pressure. Utilizing a structural equation framework, we hypothesized that: 1) The relation between age and EF is mediated by the anisotropy of white matter fibers connecting the frontal and parietal lobes, and as a control comparison, not by fibers in the corticospinal tracts, 2) This association is not driven by age-related slowing of processing speed, but 3) is influenced by the effects of white matter hyperintensity burden within frontal and parietal WM fiber tracts, and 4) differences in blood pressure serve as a salient mediating factor given its deleterious effects on white matter health.

## 2. Methods

### 2.1 Participants

The sample included 190 cognitively normal individuals recruited through flyers and media advertisements from the Dallas-Fort Worth Metroplex. Participants were sampled across the adult lifespan ranging from 20-94 years of age and were screened to be free from a history of neurological, cardiovascular, metabolic, or psychiatric problems; head trauma involving loss of consciousness > 5 minutes; substance abuse; or cognitive altering medications. Self-report screening assessments specifically excluded participants for a history of heart disease, stroke, heart attack, clinically diagnosed learning disabilities, radiation or chemotherapy treatment for cancer, HIV, contraindications to MRI such as metallic implants or claustrophobia, or anti-depressant or anti-anxiety medications. SWI scans were assessed on the sagittal, horizontal, and coronal planes and screened for micro bleeds, that of which only three participants were confirmed of having one small micro bleed. Additional inclusion requirements involved in-lab assessment of the Mini-Mental State Exam (MMSE) score > 25 (Folstein, Folstein, & McHugh, 1975), and Center for Epidemiologic Studies Depression Scale (CES-D) score ≤ 16 (Radloff, 1977), as well as thorough screening of all acquired MRI (including T1-weighted, T2-FLAIR, susceptibility weighted, diffusion weighted, and perfusion imaging) for evidence of unreported stroke or brain anomalies. Before entering the study, each participant provided written informed consent in accord with the local Institutional Review Boards. The study protocol consisted of three visits: two cognitive sessions of approximately two hours each in duration, and one MRI session approximately two hours in duration. The lag time between the two cognitive sessions was on average 1.43 weeks (SD=1.48 weeks; range = 0 - 13.68 weeks), while the lag between the second cognitive session and the MRI session was on average 7.12 weeks (SD = 5.90 weeks; range = 0 - 34.05 weeks). During quality analysis and preprocessing steps, six participants were removed from further analyses for the following reasons: low MMSE score (*n* = 1), incorrect neuroimaging data acquisition (*n* = 1), abnormalities in brain structure (*n* = 1), and inability to resolve white matter tracts of interest (*n* = 3), yielding a total *N* = 184. Demographic data for this final sample of participants are summarized in Table 1.

**Table 1:**
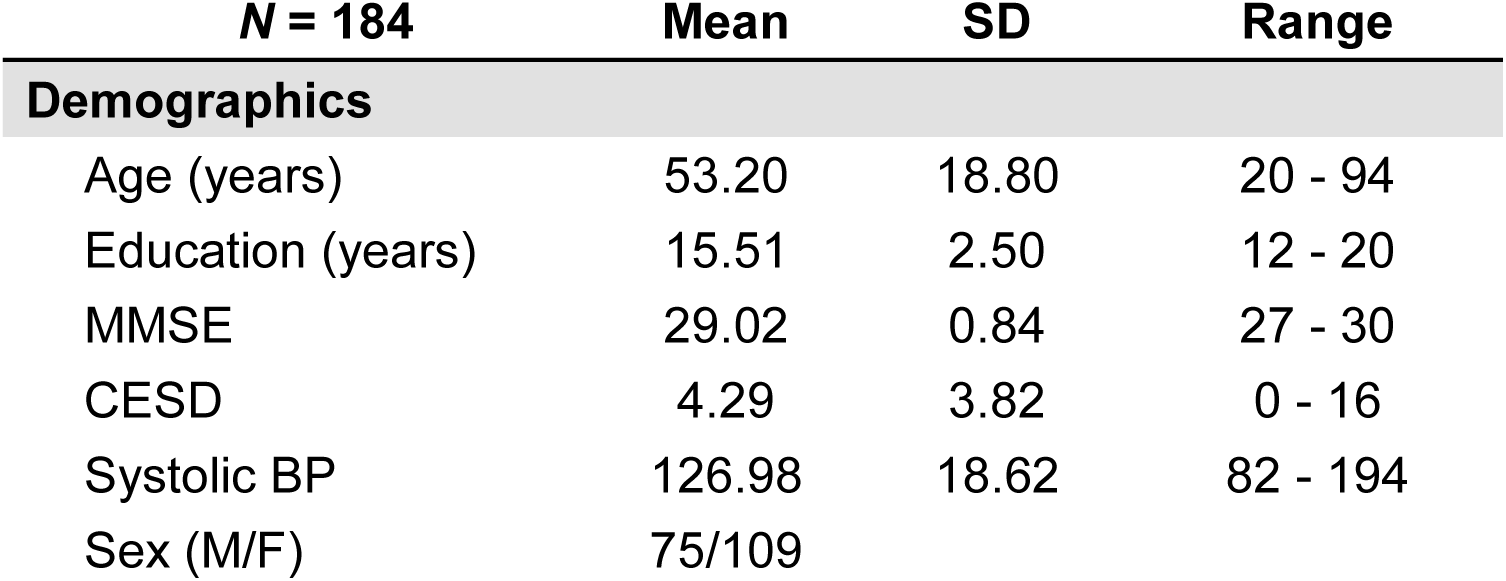
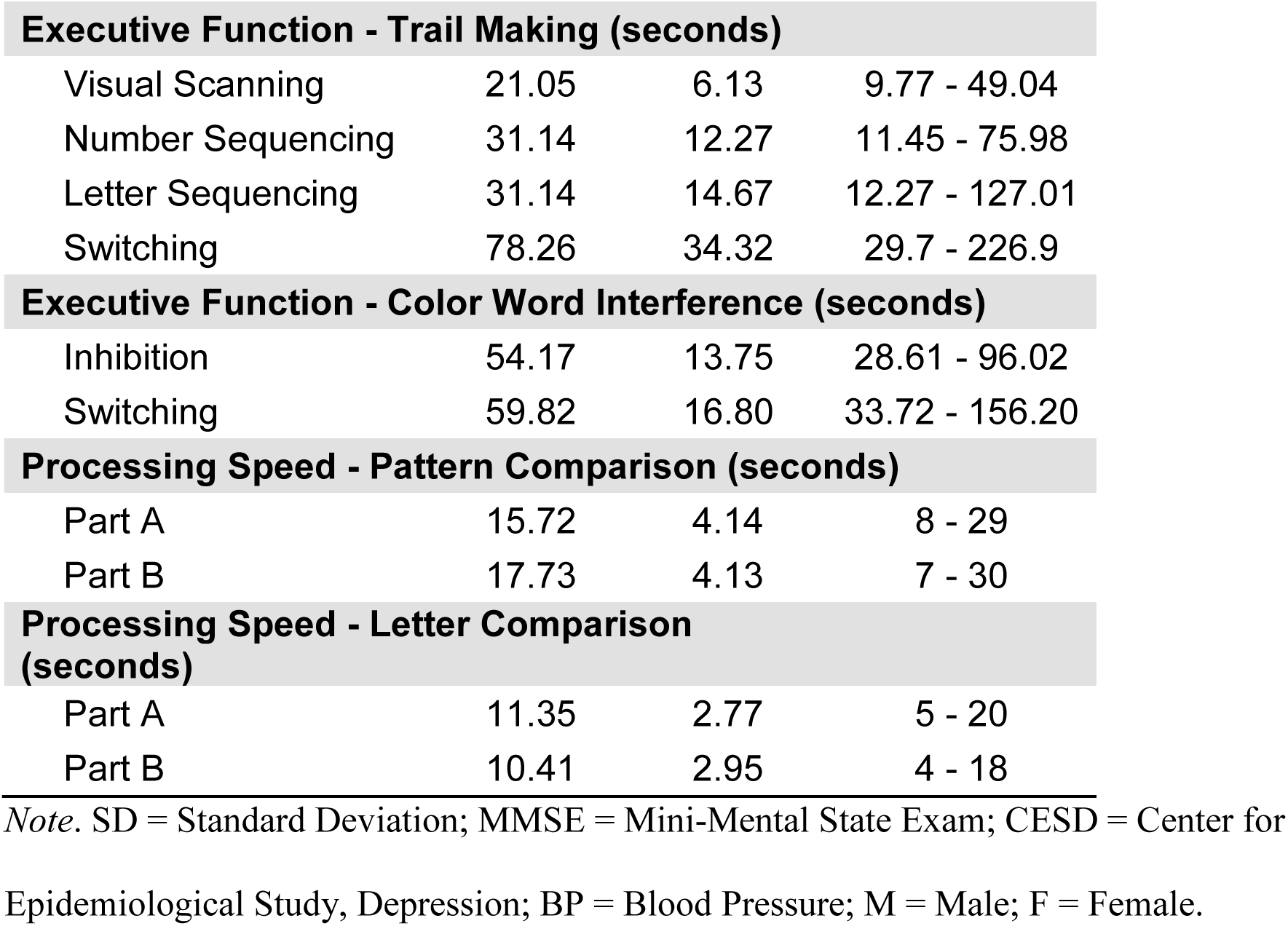
Sample demographics.

| **N = 184** | **Mean** | **SD** | **Range** |
| --- | --- | --- | --- |
| **Demographics** |  |  |  |
| Age (years) | 53.20 | 18.80 | 20 - 94 |
| Education (years) | 15.51 | 2.50 | 12 - 20 |
| MMSE | 29.02 | 0.84 | 27 - 30 |
| CESD | 4.29 | 3.82 | 0 - 16 |
| Systolic BP | 126.98 | 18.62 | 82 - 194 |
| Sex (M/F) | 75/109 |  |  |
| **Executive Function - Trail Making (seconds)** |  |  |  |
| Visual Scanning | 21.05 | 6.13 | 9.77 - 49.04 |
| Number Sequencing | 31.14 | 12.27 | 11.45 - 75.98 |
| Letter Sequencing | 31.14 | 14.67 | 12.27 - 127.01 |
| Switching | 78.26 | 34.32 | 29.7 - 226.9 |
| **Executive Function - Color Word Interference (seconds)** |  |  |  |
| Inhibition | 54.17 | 13.75 | 28.61 - 96.02 |
| Switching | 59.82 | 16.80 | 33.72 - 156.20 |
| **Processing Speed - Pattern Comparison (seconds)** |  |  |  |
| Part A | 15.72 | 4.14 | 8 - 29 |
| Part B | 17.73 | 4.13 | 7 - 30 |
| **Processing Speed - Letter Comparison (seconds)** |  |  |  |
| Part A | 11.35 | 2.77 | 5 - 20 |
| Part B | 10.41 | 2.95 | 4 - 18 |

*Note.* SD = Standard Deviation; MMSE = Mini-Mental State Exam; CESD = Center for
Epidemiological Study, Depression; BP = Blood Pressure; M = Male; F = Female.

### 2.2 Blood Pressure Measures

Participants’ blood pressure was measured at each of the three study visits using brachial cuff automatic sphygmomanometers via a Panasonic EW3153 at each cognitive session and a Welch Allyn Spot Vital Signs 420TB at the MRI session. Each measurement was taken while the participant had been in a seated position for a minimum of five minutes, arm horizontal, and legs uncrossed. Systolic and diastolic pressure were recorded in mm/Hg, as well as heart rate in beats per minute. For the current study, we utilized systolic pressure as our index of interest given evidence that systolic pressure may be a better index of cerebrovascular dysfunction than diastolic pressure (Chobanian et al., 2003) and more predictive of white matter hyperintensities (Liao et al., 1996).

### 2.3 Cognitive measures

Prior to MRI scanning, each participant underwent a comprehensive battery of cognitive testing separated across two sessions. Multiple assessment tools were used to measure executive functioning and processing speed abilities. Specifically, executive functioning was measured as completion time for each of the Color Word Interference tasks (inhibition and switching) and Trail Making subtests (visual scanning, number sequencing, letter sequencing, and switching) of the Delis-Kaplan Executive Function System (D-KEFS; Delis, Kaplan, & Kramer, 2001). Subtests that are often linked to motor and speeded abilities, such as the word reading or color naming measures in the Color Word Interference task, or the motor speed or counting 3’s assessment of the Trail Making test, were specifically excluded to avoid overlap with variance associated with processing speed. Instead, processing speed was measured using a separate task, the total number of correct responses within the 30-second duration of both parts of the Pattern Comparison and Letter Comparison tasks (Salthouse & Meinz, 1995). Descriptive statistics for cognitive scores are provided in Table 1.

### 2.4 Neuroimaging acquisition

Neuroimaging data were acquired on a single 3-Tesla Philips Achieva scanner with a 32-channel head coil using SENSE encoding (Philips Healthcare Systems, Best, Netherlands). The current study employed a T1-weighted MPRAGE high-resolution structural scan for the purposes of anatomy and registration (160 sagittal slices at a voxel size of 1mm^3^, flip angle = 12°, TR/TE/TI = 8.1ms/3.7ms/1100ms, FOV = 204×256×160, matrix = 256×256, 3:57min), a T2-FLAIR structural scan to identify areas of white matter hyperintense tissue (64 axial slices at a voxel size of .5×.5×2.5 mm^3^, flip angle=90°, TR/TE/TI = 11000ms/125ms/2800ms, FOV=230×160×230, matrix=352×212, 3:40 min), and a diffusion-weighted single-shot EPI sequence to probe white matter tracts (65 axial slices with a voxel size of 2×2×2.2 mm^3^ reconstructed to .85×.85×2.2mm^3^, 30 diffusion-weighted directions at b-value = 1000s/mm^2^ with 1 non-diffusion weighted b_0_ at 0 s/ mm^2^, TR/TE = 5608ms/51ms, FOV = 224×143×224, matrix = 112×112, 4:19min).

### 2.5 Neuroimaging data processing

#### 2.5.1 T1-weighted anatomical scans

After visual inspection for acquisition artifacts including movement distortions, high-resolution structural T1-weighted images were used as an anatomical reference, and for co-registration during multiple data processing steps, including to T2-FLAIR image space and diffusion image space, as well as between anatomical template space and diffusion space utilized in subsequent processing. Additionally, to covary for differences in head size, intracranial volume was derived by manually tracing 9 coronal slices (every 12^th^ slice) using Analyze version 12.0 (AnalyzeDirect, Overland Park, KS), summing these slices into an area, and multiplying by slice thickness to create volume in mm^3^ (as in Raz et al., 2004).

#### 2.5.2 White matter hyperintensity data

White matter hyperintense voxels were identified by processing T2-FLAIR images using the lesion prediction algorithm (LPA; Schmidt, 2017, Chapter 6.1) as implemented in the Lesion Segmentation Toolbox (LST) version 2.0.5 (www.applied-statistics.de/lst.html) for SPM12. This automated classifier produces a lesion probability map where voxel values provide an estimated probability that the corresponding T2-FLAIR voxel is a white matter lesion. Trained operators viewed each participant’s lesion probability map, overlaid on top of the participant’s T2-FLAIR image, to determine a probability threshold that best minimized false-positive voxels (e.g., motion artifacts) while leaving legitimate lesions intact. Due to image inconsistencies and variance in both the sample and the sensitivity of the LPA, probability thresholds differed among participants. False-negative voxels (e.g., the ends of periventricular “caps”) were added during the editing process to ensure that true white matter hyperintensities were included in the maps, and any false-positives that remained after thresholding were manually removed (e.g., blood vessels). Regions of avoidance (ROAs) were created to eliminate false-positive voxels in non-biologically plausible brain areas. In order to remove false-positive voxels in the ventricles (e.g., choroid plexus), each participant’s CSF probability map obtained from FMRIB’s Automated Segmentation Tool (FAST; Y. Zhang, Brady, & Smith, 2001) was registered to their native T2 space using the Advanced Normalization Tools (ANTs) software package (Avants, Tustison, & Song, 2009). The T2-registered CSF probability maps were thresholded at 0.5 and binarized to create the CSF ROA (Jenkinson, Beckmann, Behrens, Woolrich, & Smith, 2012). The remaining ROAs were placed on the Montreal Neurological Institute (MNI) 1mm space brain and registered to each participant’s native T2 space using ANTs. A mid-sagittal 2-slice thick ROA plane eliminated false-positive voxels in the septum pellucidum. A ventral ROA (below *z* = 34) removed false-positive voxels from the cerebellum and brainstem, while a dorsal ROA (above *z* = 130) removed false-positive voxels from exiting blood vessels. Whole-brain white matter hyperintensity volume maps were then binarized and registered to each participant’s native diffusion space to be used both as masks to eliminate hyperintense white matter voxels from diffusion data, and to quantify within-tract lesion load from each diffusion-based tract of interest. Additionally, as a precaution against underestimating the effects of WMHs on neighboring voxels and the associated diffusion metrics, WMH lesion maps were further dilated by a factor of 1, 2, 3, and 5 voxels to create larger, more conservative masks with which to exclude diffusion values surrounding the identified WMH. Removing these additional voxels from the various dilation masks produced no effect on the model results. Thus, all statistics, models, and figures reported hereafter use the undilated data. Illustration of WMH from a representative participant denoted by red voxels in Figure 1.

**Figure 1:**
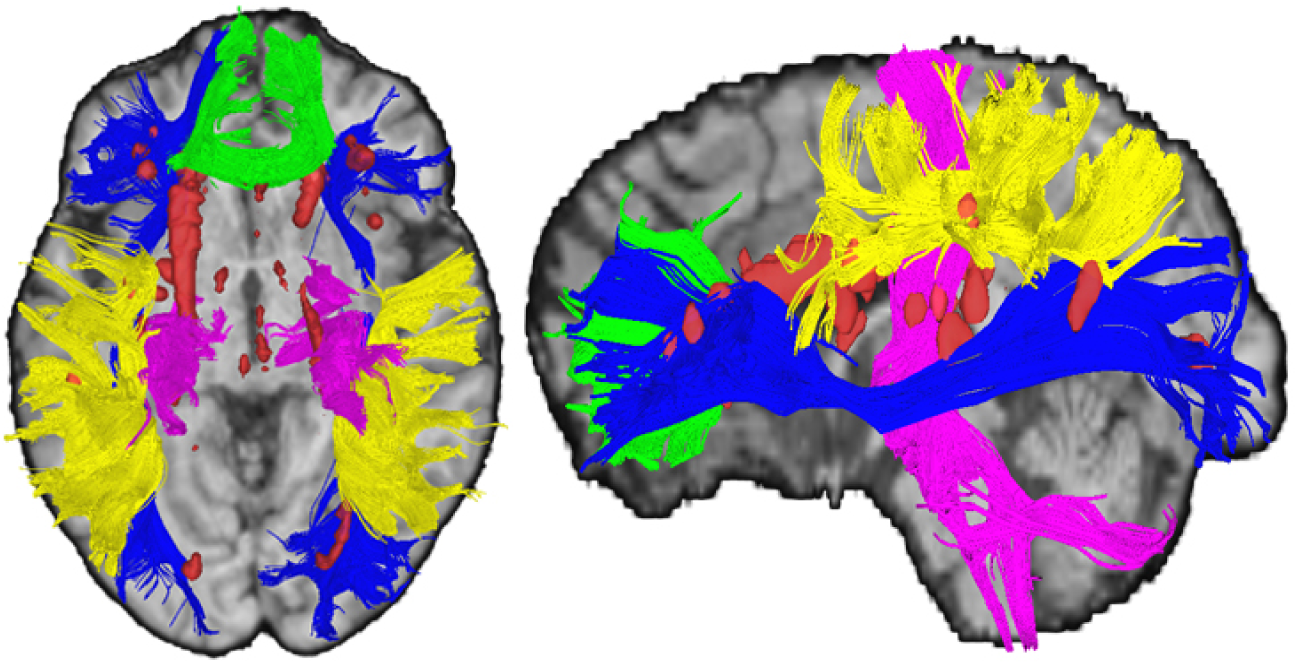
White Matter Measures. Axial and sagittal views illustrating white matter tracts resolved using deterministic tractography (green = corpus callosum genu; blue = inferior frontal occipital fasciculus; yellow = superior longitudinal fasciculus; pink = corticospinal tract) and white matter hyperintense regions (red = white matter hyperintense voxels) in a participant with an average number of white matter streamlines in each tract and both within-tract and whole brain white matter hyperintensity burden (+/- .5 SD).

#### 2.5.3 Diffusion-weighted data

Diffusion data underwent quality assurance checks utilizing both manual and automated procedures. A trained researcher identified any anatomical abnormalities or scanner artifacts across all slices of each gradient, and DTIPrep v1.2.4 (Liu et al., 2010) identified additional slice- or gradient-wise artifacts above the default thresholds. Any gradients with distortions were removed from analyses and were not incorporated into the tensor calculation. On average, less than four of the thirty gradients were identified as unusable and removed from analyses (remaining gradients ranged from 22-30). DTIPrep was then used to adjust for motion and eddy-current distortions via registration to the b_0_ image while additionally adjusting the gradient table to account for registration-induced changes to the applied encoding directions (Leemans & Jones, 2009).

Tensor calculation and deterministic tractography were conducted using DSI Studio software (build released September 26th, 2014; Yeh, Verstynen, Wang, Fernández-Miranda, & Tseng, 2013). To isolate white matter tracts of interest, a series of regions of interest (ROIs) and ROAs were strategically placed to capture representative white matter fibers while eliminating non-biologically plausible or extraneous fibers. All regions were placed on the MNI 1mm brain and registered to each participant’s native diffusion space using the ANTs software package (Avants et al., 2009). To capture white matter tracts relevant to executive functions, deterministic tractography was conducted to generate the genu of the corpus callosum (genu), the bilateral inferior frontal occipital fasciculus (IFOF), and the bilateral dorsal superior longitudinal fasciculus (SLF), and to provide a contrast projection fiber tract, the corticospinal tract (CST) was also isolated, as described below. Additional tracking parameters were applied to ensure the biological plausibility of the resolved streamlines including a fractional anisotropy (FA) threshold of .20, maximum turning angle of 60 degrees, and a minimum and maximum length of 20mm and 500mm, respectively. Tracts from a representative participant are illustrated in Figure 1.

##### CC genu

The genu was isolated using the “genu of the corpus callosum” parcellation from the JHU atlas (Mori, Wakana, Van Zijl, & Nagae-Poetscher, 2005) as an ROI. Additionally, a midsagittal ROI plane was included to ensure that resolved streamlines crossed the midline. ROAs were added to remove streamlines tracking into the cingulum (using the JHU cingulate gyrus parcellations), streamlines tracking into other corpus callosum segments (using a coronal plane posterior to the JHU genu ROI starting at slice *y* = 21), and streamlines projecting laterally (using parasagittal planes 25 slices on each side of the midline).

##### IFOF

To isolate the IFOF, two coronal 3-slice thick ROI planes were placed in the frontal (centered at *y* = 21) and occipital (centered at *y* = −75) lobes to ensure that the fiber tract traversed the entire brain. Given the extensive length of this tract, a number of ROAs are required to eliminate spurious streamlines. To eliminate streamlines more likely associated with the anterior thalamic radiations, a coronal ROA was placed between *y* = −7 and *y* = −12, but importantly did not extend axially below *z* = 3. Additionally, to remove any thalamic streamlines, in particular those that could be considered part of the internal capsule, a rectangular cuboid was placed in each hemisphere such that each stretched 38 coronal slices from *y* = 10 to *y* = −27, 17 slices axially from *z* = 7 to *z* = −9, and 5-sagittal slices between *x* = −10 and *x* = −14 in the left hemisphere and *x* = 9 and *x* = 13 in the right hemisphere. Finally, to eliminate streamlines that stray too far inferiorly (such as false frontal-temporal connections), or too far superiorly (such as frontal lobe streamlines looping towards premotor cortex), two axial ROAs planes were placed at *z* = 35 and *z* = −24.

##### SLF

The frontal-parietal limb of the SLF was isolated using two ROI coronal planes, each 12 slices thick, centered at *y* = −27.5 and *y* = −45.5. A 6-slice thick axial ROA was centered at *z* = 13.5 to ensure that the more ventral streamlines comprising the arcuate fasciculus part of the SLF were not included.

##### CST

The projection fibers of the CST were isolated by using two ROIs and four ROAs. The ROIs consisted of the precentral gyrus, delineated from within the FreeSurfer software package using the Desikan-Killiany atlas (Desikan et al., 2006), and the brainstem, defined by a region that stretched axially from *z* = −15 to *z* = −50, sagittally from *x* = −15 to *x* = 17, and coronally from *y* = −13 to *y* = −36. ROAs consisted of a 3-slice thick coronal plane (centered at *y* = 21), two 5-slice thick sagittal exclusion planes at x = −43 and x = 42, the lateral occipital region from the Desikan-Killany atlas, and a 3-slice thick midsagittal plane to eliminate fibers crossing the midline.

Visual inspection ensured that each tract was accurately resolved for each participant based upon previous literature and anatomical atlases (Catani & De Schotten, 2008; Huang et al., 2005; Makris et al., 2005). Three participants were removed at this stage due to the inability to sufficiently resolve the IFOF (less than 100 streamlines resolved). Each participant, on average, had 4,454 (± 1187 SD) streamlines in the genu, 6048 (± 1946 SD) streamlines in the SLF, and 3137 (± 1348 SD) streamlines in the IFOF. These streamlines served as data-driven regions of inclusion to identify the voxels most likely contributing to structural connections between the frontal and parietal regions. Lesion maps were used as masks to remove white matter hyperintense voxels from those contained within the resolved tracts to avoid inclusion in any calculation quantifying white matter diffusion. FA values from the remaining voxels contained within each of the streamlines were averaged across tract and across hemisphere (for SLF and IFOF) to obtain a single FA value for each of the three white matter tracts.

### 2.6 Data analysis and model specification

To investigate the complex associations among age, blood pressure, brain, and cognitive variables, data were analyzed using structural equation modeling (SEM) within Mplus v8 statistical software (Muthén & Muthén, 2017) using maximum likelihood estimation and 5000 bootstrap iterations. Measurements for which there was a single value collected, such as age, and covariates of years of education, sex, and intracranial volume, were modeled as observed/manifest variables. All other measures were used as observed indicators to form latent variables to capitalize on covariance among data points. SEM allows for variables with a high degree of covariance to co-exist and account for one another within the same structure, thus allowing for assessment of the unique variance attributable to each variable. Given the known overlap in variance shared between EF and PS (Salthouse, 2011), SEM is essential to allow for modeling of the unique cognitive contributions of each process. Cognitive measures were combined into two latent variables, one for EF and one for PS (see measurement model fit indices in Supplemental Information), by standardizing each of the individual tasks into *z*-scores and modeling them together within their associated domain. White matter hyperintensity data were modeled by individually *z*-scoring the within-tract lesion loads (volume in mm^3^) from the genu, SLF, and IFOF voxels and allowing them to covary as a single white matter hyperintensity latent variable (denoted as WMH in subsequent text and models). Similarly, fractional anisotropy was modeled by individually *z*-scoring the FA values from each tract and allowing them to covary as a single white matter fractional anisotropy latent variable (denoted as WMFA in subsequent text and models). Blood pressure measures, collected at each time point during the study (cognitive session one, cognitive session two, and MRI), were combined into a latent variable using the systolic measurements from each data point (denoted as BP in subsequent text and models). All participants had a complete data set with the exception of a few individuals missing BP readings (maximum of 4 data points) due to equipment malfunction. In addition to using 95% CI with 5000 bootstraps, to acknowledge potential inflation of familywise error buildup across estimated paths, 99% CI were also estimated. A correlation matrix of all variables modeled is included in Table 2.

**Table 2:**
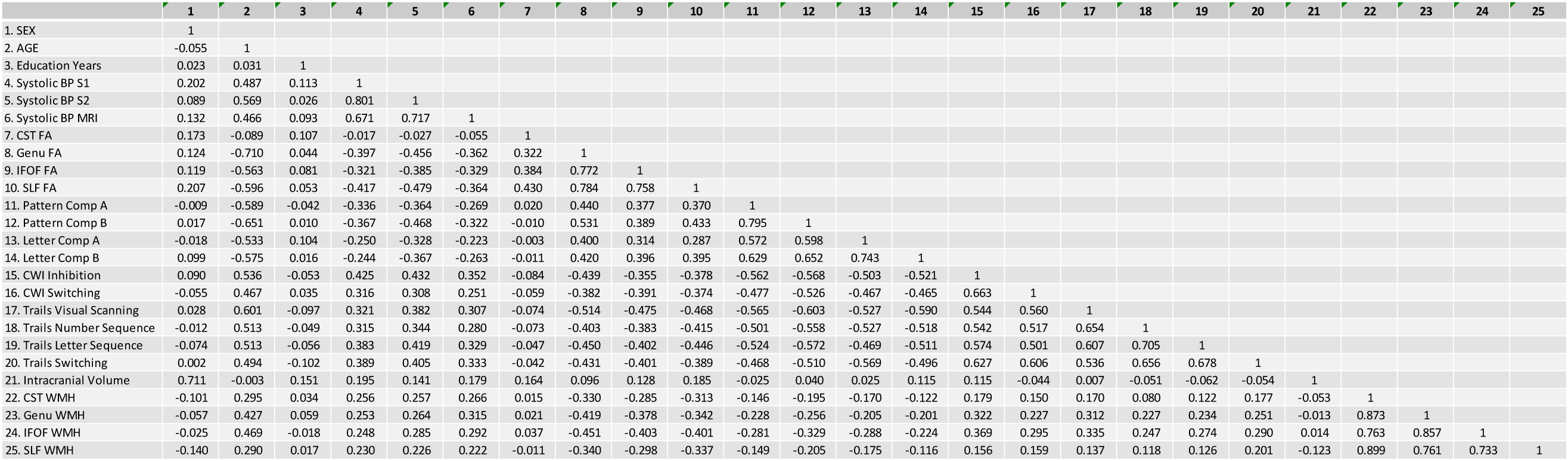
Correlation matrix of modeled variables.

To address the hypotheses, four separate structural equation models were created. Each model was designed to incrementally build upon the previous models in order to gain more insight into the mediating effects of brain- and health-related variables on the association between age and executive functioning. Model fit was estimated by examining the following fit indices: Akaike Information Criterion (AIC), Bayesian Information Criterion (BIC), Root Mean Square Error of Approximation (RMSEA; < 0.01 indicates excellent, < 0.05 good, < 0.08 moderate fit to the data), Comparative Fit Index (CFI) and Tucker-Lewis Index (TLI; excellent fit ≥ 0.95), Standardized Root Mean-Square Residual (SRMR; excellent fit ≤ 0.05, good fit < 0.08), and Chi-Square Test of Model Fit for Baseline Model (*p*-value) (Raykov & Marcoulides, 2006). In lieu of traditional power analysis, the sample size for structural equation modeling followed the rule of thumb of greater than 10 participants per path estimated (e.g., for our most complex model, 10 paths were estimated, requiring at least 100 participants). Data and code for this study are publicly available via OSF at https://osf.io/hts8w/?view_only=9c8ce88227cf47c79a1d3e11dd0d6363. No portion of the study procedures or analysis was preregistered. All participant and data inclusions and exclusions were determined before study analyses were conducted.

## 3. Results

Each model demonstrated acceptable fit parameters as detailed in Table 3. In each model diagram (Figures 2-5), squares represent observed variables, circles represent latent variables, straight dark lines are significant path estimates, curved lines are autocorrelations, thin lines represent factor loadings, and light gray lines indicate non-significant paths. Values along the paths are standardized beta estimates with 95% CIs (significant when the range does not include zero), showing direction and strength of the effect. As a penalty for multiple estimations, 99% CIs were also estimated. All significant paths at 95%CI remained significant at 99%CI (i.e., did not include 0) in all models.

**Figure 2:**
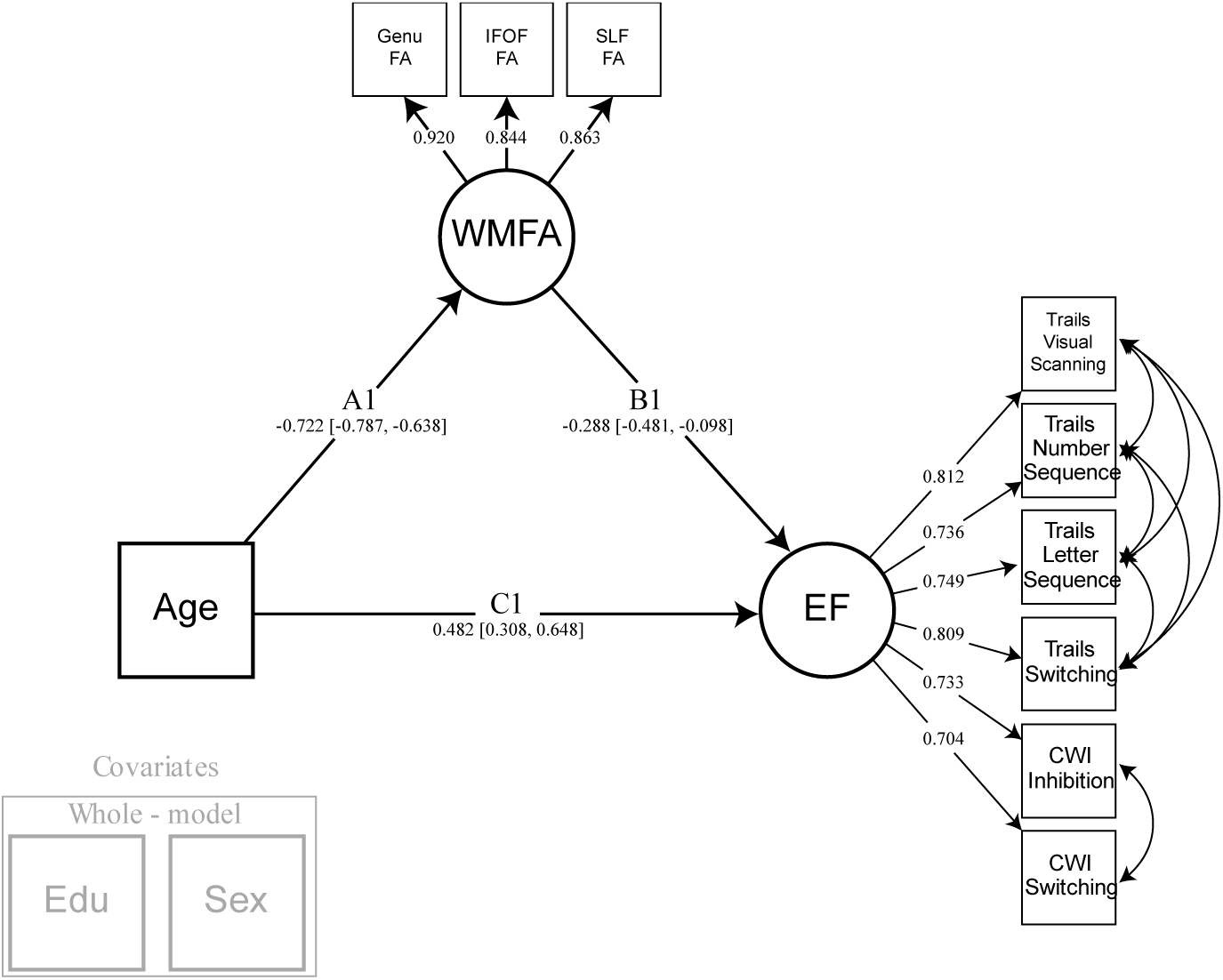
Model 1. Simple mediation model of age by executive function, with white matter fractional anisotropy of the fronto-parietal white matter tracts. Years of education and participant sex were regressed from all latent variables modeled. Note: EF = executive function; WMFA = white matter fractional anisotropy; FA = fractional anisotropy; IFOF = inferior frontal occipital fasciculus; SLF = superior longitudinal fasciculus; CWI = color word interference; EDU = years of education.

**Table 3:** Summary of structural equation model fit metrics.

| **Model Fit Parameter** | **Model 1** | **Model 2** | **Model 3** | **Model 4** |
| --- | --- | --- | --- | --- |
| Akaike Information Criterion (AIC) | 3530.68 | 5016.41 | 6148.79 | 7305.43 |
| Bayesian Information Criterion (BIC) | 3662.49 | 5209.31 | 6402.77 | 7604.42 |
| Root Mean Square Error of Approximation (RMSEA) | 0.049 | 0.047 | 0.047 | 0.044 |
| Comparative Fit Index (CFI) | 0.986 | 0.982 | 0.977 | 0.976 |
| Tucker-Lewis Index (TLI) | 0.977 | 0.974 | 0.969 | 0.969 |
| Standardized Root Mean Square Residual (SRMSR) | 0.028 | 0.03 | 0.037 | 0.04 |
| Chi-Square Test of Model Fit for Baseline Model (*p*-value) | < 0.001 | < 0.001 | < 0.001 | < 0.001 |

### 3.1 Mediating effect of frontal-parietal white matter on the age-EF relationship (Model 1)

The first goal of this study was to determine whether the proxies of white matter health, accessed at the higher-order cognitive centers of the frontal and parietal regions, played a significant role in the age-related decline in executive function abilities commonly observed with aging. This simple mediation model (Figure 2) was specified to test a direct effect of age on the EF latent construct, and an indirect effect where the WMFA latent construct mediates the age/EF association, while covarying for education and sex. Significant standardized path estimates were found for the association between age and WMFA (Path A: Est. = −.722, 95% CI [-.787, −.638]), WMFA and EF (Path B: Est. = −.288, 95% CI [-.481, -.098]), and age and EF (Path C: Est. = .482, 95% CI [.308, .648]). Given that higher EF values indicate poorer performance, and lower FA values indicate less structural impedance and directionality in white matter tissue, these results show main effects of each association in the expected direction. Importantly, a significant indirect effect was found for the mediation of WMFA on the relationship between age and EF (Path A*B: Est. = .208, 95% CI [.071, .356]).

### 3.2 Mediating effect of frontal-parietal white matter on the age-EF relationship accounting for PS (Model 2)

After confirming that frontal-parietal white matter FA mediates the association between age and executive function, we sought to determine the functional specificity of these white matter connections by introducing PS as an outcome variable. Model 2 was created by adding a PS latent variable to the Model 1 specification as a separate cognitive construct for which WMFA could be mediating (Figure 3). Both cognitive constructs were allowed to covary to account for any shared variance among the individual measures and to control any potential overlap between these cognitive domains. For measurement model fit indices for the cognitive variables, see Supplemental Information. Significant standardized path estimates were found for the association between age and WMFA (Path A: Est. = -.664, 95% CI [-.770, -.545]), WMFA and EF (Path B1: Est. = -.253, 95% CI [-.467, -.085]), age and EF (Path C1: Est. = .398, 95% CI [.268, .522]), and between age and PS (Path C2: Est. = -.534,95% CI [-.697, -.383]). The path between WMFA and PS, notably, was not significant. A significant indirect effect was found for the mediation of WMFA on the association between age and EF (Path A*B1: Est. = .208, 95% CI [.076, .352], but there was no mediation found for age and PS, suggesting some specificity of these associations to executive functioning.

**Figure 3:**
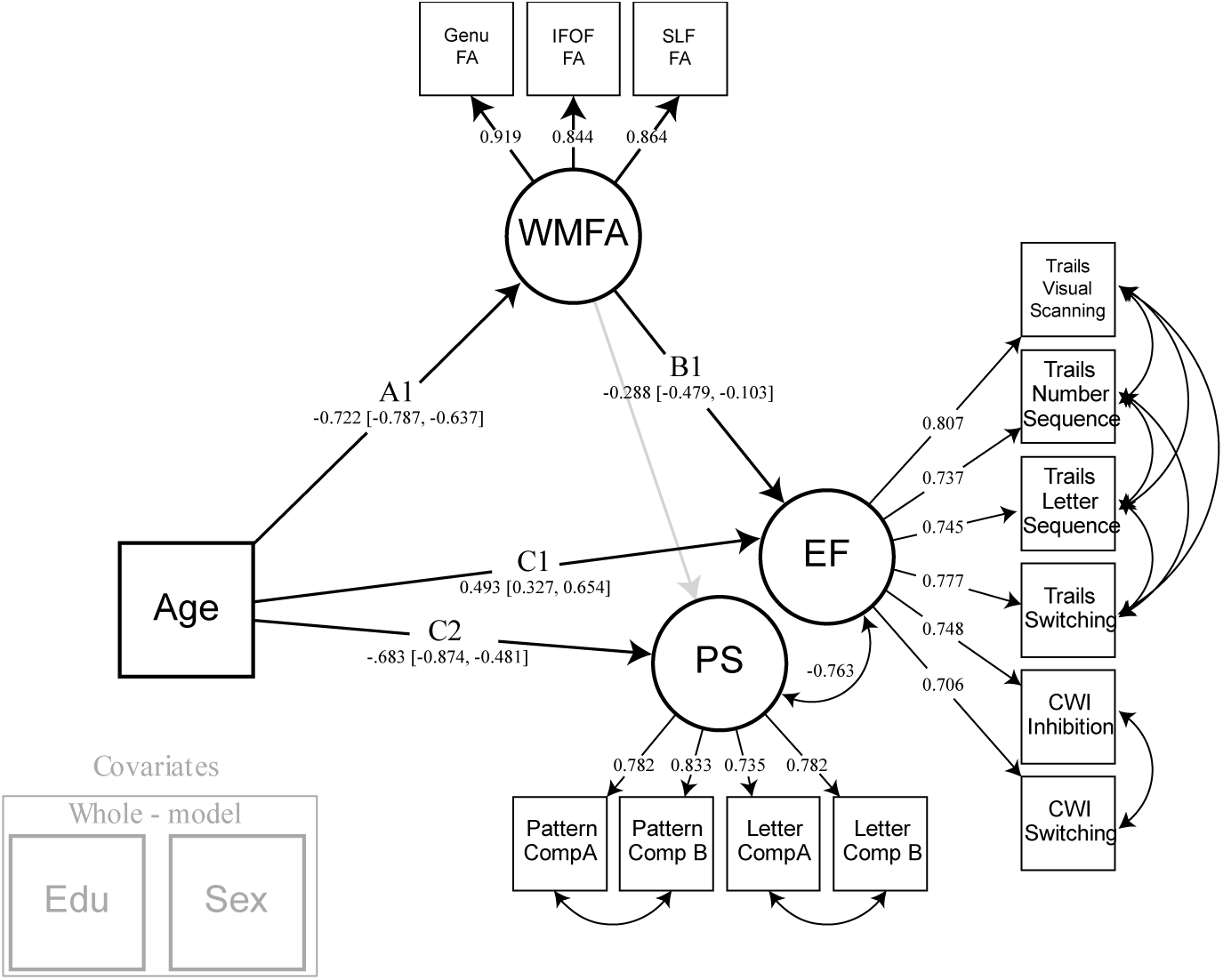
Model 2. Model 2 builds upon model 1 by adding processing speed as a cognitive construct. Processing speed serves to regress speed-like features from the executive function construct to demonstrate specificity of the selected white matter tracts in their association to switching and inhibition. Years of education and sex were regressed from all latent variables modeled. Note: EF = executive function; PS = processing speed; WMFA = white matter fractional anisotropy; FA = fractional anisotropy; IFOF = inferior frontal occipital fasciculus; SLF = superior longitudinal fasciculus; CWI = color word interference; EDU = years of education.

In addition to the observed association between white matter FA and EF but not PS, we also sought to demonstrate that this effect was specific to the white matter connecting frontal and parietal regions. Using the CST white matter as a comparison region, FA values were added as an additional observed variable (CSTFA) that could be mediating the age-PS or age-EF relationship, while allowing for covariance between both white matter latent variables (Model 2A, Figure 4). Significant standardized path estimates were found for the association between age and WMFA (Path A1: Est. = -.717, 95% CI [-.783, -.631]), WMFA and EF (Path B1: Est. = -.372, 95% CI [-.605, -.141]), age and EF (Path C1: Est. = .444, 95% CI [.262, .622]), and between age and PS (Path C2: Est. = -.620, 95% CI [-.826, -.401]). Interestingly, the path between WMFA and PS remained non-significant, and the path between CSTFA and EF was non-significant. In contrast, the path between CSTFA and PS was significant (*p*=0.035), however the 95% confidence interval includes 0 after 5000 bootstraps (Path EB1: Est. = −0.150, 95% CI [-.320, .013]. While this association between CSTFA and PS is weak at best, it demonstrates that the relationship between white matter and our executive function latent is unique to the frontal-parietal latent, and that other white matter tracts, such as the CST, could be related to other aspects of cognition. Note, that CST is modeled as a single indicator rather than a latent factor, and as such differs in its ability to parse error variance compared to latent constructs such as the WMFA.

**Figure 4:**
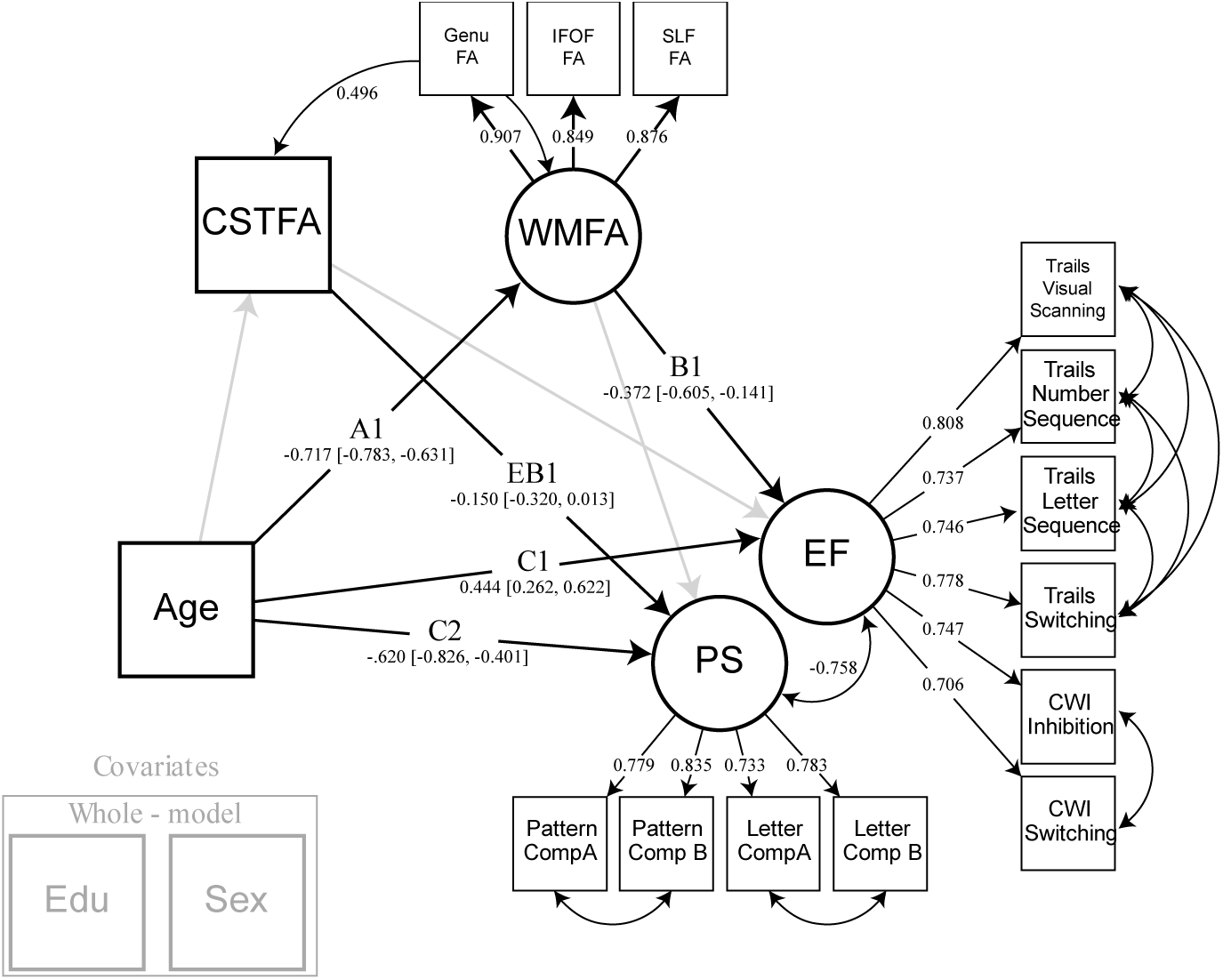
Model 2A. Model 2A demonstrates the specificity of the frontal-parietal white matter by adding an additional white matter pathway, the corticospinal tract, to the model. The CST not only allows for covariance between two distinct white matter regions, but it also shows that different white matter pathways are related to separate aspects of cognition. Years of education and sex were regressed from all latent variables modeled. Note: EF = executive function; PS = processing speed; CSTFA= corticospinal tract fractional anisotropy; WMFA = white matter fractional anisotropy; FA = fractional anisotropy; IFOF = inferior frontal occipital fasciculus; SLF = superior longitudinal fasciculus; CWI = color word interference; EDU = years of education.

### 3.3 Influence of white matter hyperintensities on white matter tract structure and cognition (Model 3)

After establishing the specificity of these white matter tracts in their relationship with EF, and accounting for shared variance between the two cognitive constructs, we examined the potential deleterious effects of white matter hyperintensities on the structure of white matter fibers. Prior literature indicates that white matter hyperintensity burden increases with age and influences the integrity of white matter fiber pathways, leading to declines in cognitive performance. Therefore, Model 3 (Figure 5) was specified to build off of Model 2 by adding paths that reflect the influence of WMH on WMFA, while also considering the separate mediating influences WMH may contribute directly to cognitive processing. Additionally, intracranial volume was added as a covariate to account for the volumetric nature of measuring white matter hyperintensity burden. Significant standardized path estimates were found for the following associations: age and WMFA (Path A1: Est. -.638 =, 95% CI [-.722, -.538]); age and WMH (Path A2: Est. = .461, 95% CI [.354, .550]); WMFA and EF (Path B1: Est. = -.272, 95% CI [-.481, -.090]); WMH and WMFA (Path D1: Est. = -.189, 95% CI [-.275, -.095]); age and EF (Path C1: Est. = .490, 95% CI [.309, .674]); and age and PS (Path C2: Est. = -.703, 95% CI [-.905, -.502]). The paths between WMH and each of the cognitive constructs were not significant, nor was the relation between WMFA and PS. A significant indirect effect was found, however, for the mediating path that represented WMH and WMFA’s influence on the association between age and EF (Path A2*D1*B1: Est. = .019, 95% CI [.004, .048]). This indicates that WMH burden influences WMFA and that together this path significantly mediates the age-EF relationship. However, this was specific to EF as there remained no significant mediating effect on age and PS. Additionally, the original mediation of WMFA on the age and EF relationship remained significant after accounting for the effects of WMH (Path A1*B1: Est. = .140, 95% CI [.045, .265]). Finally, to test a separate model that examines any possible influence of WMFA on WMH, we reversed the WMH and WMFA path direction and found no significant mediation in this direction (data not shown).

**Figure 5:**
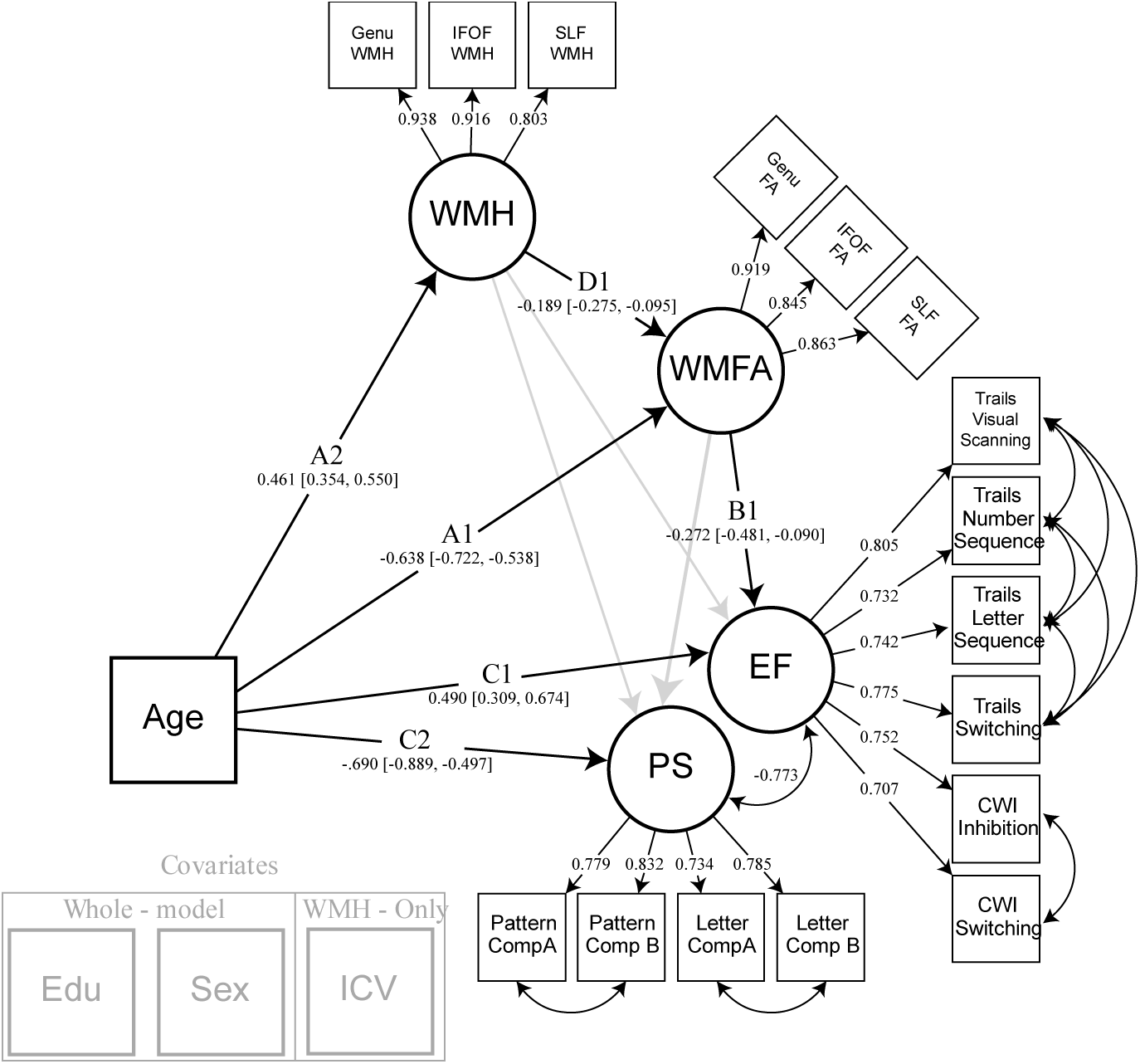
Model 3. Model 3 builds upon model 2 by adding within-tract white matter hyperintensity burden as a latent variable that directly influences white matter fractional anisotropy. This allows examination of the detrimental effects of lesion-like infarcts within white matter tracts of interest, and their overall influence on cognitive abilities. Years of education and sex were regressed from all latent variables modeled. Intracranial volume was regressed from white matter hyperintensity burden. Note: EF = executive function; CWI = color word interference; PS = processing speed; WMFA = white matter fractional anisotropy; FA = fractional anisotropy; IFOF = inferior frontal occipital fasciculus; SLF = superior longitudinal fasciculus; WMH = white matter hyperintensity burden; EDU = years of education; ICV = intracranial volume.

### 3.4 Influence of blood pressure on WMH and WMFA leading to poorer cognitive performance (Model 4)

To assess the influence of individual differences in vascular health on frontal-parietal white matter FA and within-tract white matter hyperintensity burden, we incorporated an estimate of blood pressure in our model. A latent variable composed of three systolic blood pressure measures was added to the model as a mediator of the relationship between age and WMH, as well as age and WMFA (Figure 6). This also indirectly influences the mediation of the associations between age and cognition, though we hypothesized that this would occur via a direct influence on white matter factors, but not be related to cognition itself. Significant standardized path estimates were found for the associations between age and WMFA (Path A1: Est. = -.525, 95% CI [-.664, -.373]), age and WMH (Path A2: Est. = .407, 95% CI [.285, .577]), age and BP (Path A3: Est. = .597, 95% CI [.483, .686]), WMFA and EF (Path B1: Est. = -.283, 95% CI [-.496, -.096]), WMH and WMFA (Path D1: Est. = -.175, 95% CI [-.260, -.088]), BP and WMFA (Path D3: Est. = -.196, 95% CI [-.364, -.044]), age and EF (Path C1: Est. = .483, 95% CI [.299, .670]), and age and PS (Path C2: Est. = -.701, 95% CI [-.902, -.503]). The relation between WMH and each of the cognitive constructs, as well as the relation between WMFA and PS, remained non-significant. Additionally, the relation between BP and WMH was also non-significant. The indirect effect previously found for the mediating path that represented WMH and WMFA’s influence on the relationship between age and EF (Path A2*D1*B1: Est. = .020, 95% CI [.005, .047], and WMFA alone on age and EF (Path A1*B1: Est. = .149, 95% CI [.059, .268]), both remained significant. The new mediation path that examined the combined effects of BP and WMFA on the relation of age and EF was also significant (Path A3*D3*B3: Est. = .033, 95% CI [.005, .097]). However, any path including both BP and WMH was non-significant.

**Figure 6:**
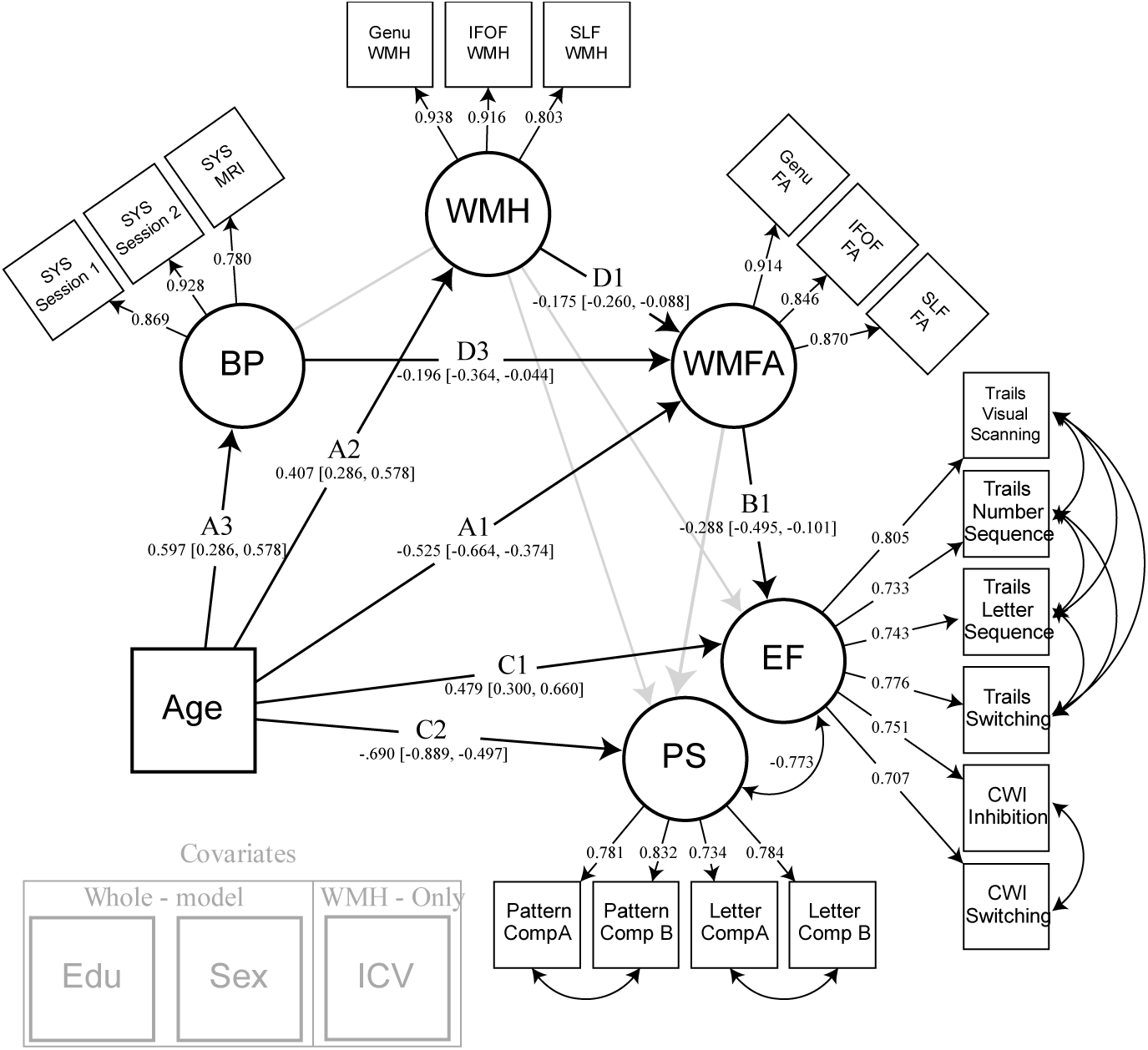
Model 4. Model 4 builds upon model 3 by adding systolic blood pressure as a latent variable that directly influences both white matter hyperintensity burden and white matter fractional anisotropy. Years of education and sex were regressed from all latent variables modeled. Intracranial volume was regressed from white matter hyperintensity burden. Note: EF = executive function; CWI = color word interference; PS = processing speed; WMFA = white matter fractional anisotropy; FA = fractional anisotropy; IFOF = inferior frontal occipital fasciculus; SLF = superior longitudinal fasciculus; WMH = white matter hyperintensity burden; BP = blood pressure; SYS = systolic; EDU = years of education; ICV = intracranial volume.

## 4. Discussion

The present study identified key factors that influence age-related differences in cognitive performance by identifying specific contributions of tract-specific white matter anisotropy, white matter hyperintensity burden, and blood pressure through an incremental structural equation modeling approach. Executive function performance is highly vulnerable to the effects of aging, and is worse in participants who demonstrate poorer diffusion properties of fronto-parietal (but not cortico-spinal) white matter. More specifically, decreased fronto-parietal fractional anisotropy mediates executive function abilities with age, while additionally, increased blood pressure and white matter hyperintensities within these white matter tracts each contribute to poorer white matter health and are important contributors to overall brain health.

The primary finding, that the relationship between age and EF is mediated by the health of fronto-parietal white matter connections, was motivated by the growing body of literature emphasizing the importance of network-like brain organization between higher-order association cortices that is responsible for efficient cognitive processing in aging (Damoiseaux, 2017; Fjell et al., 2016). These results advance both the idea of regional vulnerabilities in the aging brain and the disconnected brain theory of cognitive aging. Regional vulnerabilities in tissue structure have been reported across imaging modalities with an emphasis on the higher-order cognitive centers typically found in the frontal and parietal cortices (Hoagey, Rieck, Rodrigue, & Kennedy, 2019), especially with regard to executive function performance (Bennett & Madden, 2014; Bettcher et al., 2016; Cole et al., 2013; Kennedy & Raz, 2009a; Llufriu et al., 2017). The observed specificity of the white matter interconnecting frontal and parietal cortices, but not of the projection fibers of the CST, adds support to theories of retrogenesis or last-in, first-out views of age-related decline, while demonstrating the cognitive correlates of these differences in structural integrity. Decreases in fractional anisotropy point to a lessening in the overall directionality of water flow along these fronto-parietal tracts, suggesting a number of failures could be at play such as decreased or thinning of the myelin sheath, loss of fiber organization, axonal damage, or decreased density or thinning of fiber tracts (Bartzokis, 2004; Davis et al., 2009; Madden et al., 2009). Future research is needed to examine possible microstructural changes at the neuronal level (i.e., biophysical diffusion models). Notably, in a separate class of white matter, the corticospinal tract, thought to mature early in life to enhance motor responses during development, there was no evidence of an age-related vulnerability. While this does not eliminate the possibility that additional higher-order white matter pathways are also related to executive function performance, it does provide specificity that speaks toward the retrogenesis hypothesis.

Regardless of the mechanisms leading to poorer anisotropy, these results point to a disconnection of the communication among higher-order cognitive cortices, which does not appear to be a ubiquitous, brain-wide consequence of differences in white matter health. Cortical disconnections result in a loss in communication efficiency of neurons and are evidenced by associations between cognitive declines and metrics demonstrating impairment in the health of functional and structural connectivity (Cox et al., 2016; Langen et al., 2018). With regional specificity, we are able to demonstrate that measures indicative of structural connectivity are, at least partially, related to the efficiency in EF performance. Identifying and understanding the specificity involved in the relationship between the brain and cognitive abilities is critical for localizing vulnerabilities and could lead to biomarker-like targets, for both clinical and aging research, and help develop strategies for preserving cognitive abilities (Madden et al., 2017).

Another goal of the study was to interrogate the specific white matter connections of the genu of the corpus callosum, IFOF, and SLF, to evaluate their role in cognitive specificity, compared to a projection fiber tract not expected to be directly involved in EF performance. We found that differences in white matter anisotropy in these fronto-parietal tracts are related to executive function performance, but not to processing speed. The EF construct modeled in this study taps several EF subcomponents, as it is composed of switching and inhibition measures, but might not generalize to other EF subdomains (such as temporal ordering or attention). However, importantly, we believe this construct to be independent from the influence of processing speed given the covariance with speed-based simple comparison tasks. This approach allowed for an avoidance of overlap that might exist between the cognitive tasks, and instead, constructs were modeled as residualized versions of each other. As hypothesized, this method revealed no significant results related to processing speed, likely due to the specificity involved in selecting white matter tracts mirroring those pathways active during EF processing. These results demonstrate an independent contribution of fronto-parietal white matter connectivity that stands in contrast to previous work linking processing speed to more general aspects of white matter or age-related slowing (Penke et al., 2010), or neurocognitive test batteries with a high cognitive burden unaccounted for in processing speed tasks (Kerchner et al., 2012; Salami, Eriksson, Nilsson, & Nyberg, 2012).

Vascular risk factors have been proposed as a candidate mechanism for the disruptions in white matter health leading to declines in cognitive performance via increases in white matter hyperintensity burden or increased blood pressure (Bender & Raz, 2015; de Groot et al., 2001; Jacobs et al., 2013; Kennedy & Raz, 2009a; Langen et al., 2018; Langen et al., 2017; Power et al., 2017; Salat et al., 2012). In the current study, the influence of white matter hyperintensity burden is modeled directly and related to fractional anisotropy of the white matter, contributing to the mediation of age and EF, but not directly related to cognitive performance. While this builds on traditional “lesion” models with the inclusion of within-tract white matter hyperintensity burden, the findings differ from what would be expected in that these vascular lesions are not directly related to cognitive performance, but rather seem to only show a relation to proxies of white matter health. To confirm this outcome, we reversed the WMH and WMFA constructs path direction in model 4 to verify the directionality of the association and found that WMFA was indeed, unrelated to WMH and that the overall mediation was no longer significant, demonstrating additional specificity inherent to the model. Much discussion has been raised around the concerns of WMH contamination of diffusion imaging metrics such as FA (e.g., Leritz et al., 2014; Pelletier et al., 2016; Svärd et al., 2017; Van Leijsen et al., 2018; Vangberg, Eikenes, & Håberg, 2019; Vernooij et al., 2008), and the current study provides some information to that discussion, namely that WMFA is a salient predictor of age-related cognitive performance differences, independent of any effect of WMH lesions (measured at several levels of WMH lesion dilation kernel sizes).

Additionally, while systolic blood pressure measures were related to white matter FA and contributed to the mediation of age and executive function performance, no direct association of BP with white matter hyperintensity burden was detected. Therefore, it would appear that increased blood pressure and white matter hyperintense areas may separately influence white matter health, and that the cross-sectional association of blood pressure and WMH burden is weak in the current sample. This finding appears to stand in contrast to much of the literature, but may be due to our sample selection of highly healthy adults with relatively mild increases in blood pressure, less pronounced white matter hyperintensity burden, and exclusion of individuals with clinical concerns such as small vessel disease. Previous research groups have also reported that this association is weak in healthy participants with only mild vascular risk (Lindemer, Greve, Fischl, Augustinack, & Salat, 2017), or is only found with more clinically advanced white-matter degeneration, and not present in samples including younger individuals (Maillard et al., 2012). However, the idea of separate mechanistic influences of blood pressure and white matter hyperintensity influence on age-related cognitive decline has previously been discussed (Jacobs et al., 2013). The current results suggest that these vascular risk factors might exert varying effects on cognitive performance because of their unique influences on different properties of white matter integrity. It has also been posited that ontogenetically and phylogenetically later regions are differentially associated with vascular risk compared to earlier developing regions, perhaps following an anterior-posterior gradient. This differential pattern follows a retrogenesis theory, where healthy, uncomplicated aging is associated with anterior white matter changes (i.e., frontal association regions), but the addition of vascular risk pushes an expansion to more posterior regions (i.e., parietal association cortices), both in diffusion-based studies (Kennedy & Raz, 2009) and in studies of longitudinal WMH progression over 5 years (Raz et al, 2007). Specific work remains to be done to test this interesting hypothesis. Overall, these findings highlight the significant role that vascular factors play in brain aging processes, which is predictive of age-related cognitive decreases even within healthy adults.

### 4.1 Considerations and Conclusions

The current findings should be considered in the context of their strengths and limitations. The methods utilized in the current project offer several advantages over much of the current literature. Our use of diffusion tractography allowed an extraction of diffusivity estimates from the entirety of each white matter tract at the voxel level, as opposed to using a skeletonized version derived from a group template. These tracts were uniquely generated in each individual, included a “control” region, and allowed for within-tract quantification of white matter hyperintensity burden and removal of white matter hyperintense voxels from fractional anisotropy calculations, with several dilation iterations. Additionally, the use of structural equation modeling allowed for all of the variables of interest to be modeled simultaneously in such a way that they account for possible instances of covariance. The complexity inherent to model our hypotheses of interest required that we use a method that accounts for shared variance among multiple cognitive constructs, various measures of white matter health, and vascular factors, while removing additional confounds such as intracranial volume, sex, and education. Without appropriate modeling, the study results would lack specificity and instead point to more generalized correlations among white matter variables or broad cognitive constructs as opposed to showing specificity with executive function performance (Salthouse et al., 2015). While specificity is a strength of this current project, it limits the scope of our findings such that other research examining a broader range of white matter and/or cognitive tasks may find that other white matter tracts are specifically related to processing speed or to additional aspects of EF not measured in the current study (Borghesani et al., 2013). An additional caveat is that structural equation modeling is a regression-based technique and is unable to test causality or test lead-lag relationships in data collected in a cross-sectional fashion. Further, these models are only a test of one set of *a priori* configured hypotheses; other variables and modeling configurations could be used to test related hypotheses. Nevertheless, the present results highlight the importance of vascular health on the efficient communication of fronto-parietal white matter and the negative impact this has on executive function performance in aging. Additional longitudinal analyses to fully establish possible lead-lag relationships among these variables are currently underway and should soon shed light on how health factors lead to structural change and cognitive decline.

## Acknowledgements

This study was funded in part by grants R00-AG036848, R00-AG-036818, R01-AG-056535 from the National Institute on Aging. We thank Samantha Owens for assistance with white matter hyperintensity measurements and visual inspection. We report how we determined our sample size, all data exclusions, all inclusion/exclusion criteria, whether inclusion/exclusion criteria were established prior to data analysis, all manipulations, and all measures in the study.

## Supplemental Information

**Supplemental Table S1.** Cognitive variable factor structure. Measurement models of all cognitive variables of interest as either a one-factor solution or as a two-factor solution (executive function tests and processing speed tests as separate constructs). The 2-factor solution fit the data significantly better than a 1-factor solution (via a chi-square difference test *χ2*[9] = 103.006, *p* < .00001.)

| Model Fit Parameters | 1-Factor | 2-Factor |
| --- | --- | --- |
| **Akaike Information Criterion (AIC)** | **4208.56** | **4123.554** |
| **Bayesian Information Criterion (BIC)** | **4305.008** | **4248.936** |
| **Root Mean Square Error** | **0.162** | **0.126** |
| **Comparative Fit Index (CFI)** | **0.861** | **0.938** |
| **Tucker-Lewis Index (TLI)** | **0.822** | **0.893** |
| **Standardized Root Mean Square Residual** | **0.058** | **0.038** |
| **Chi-Square Test of Model Fit** | **204.463** | **101.457** |

**Supplemental Table S2.** Alternative model 2 fit parameters. Two separate specifications of the original “Model 2” were tested to compare the fit parameters when executive function and processing speed were separated and covarying with each other (EF/PS) and for when these measures were modeled as a single cognitive latent (COG). While both models demonstrate excellent fit parameters, the EF/PS specification demonstrated a slight improvement over the COG model in many of the most commonly referenced fit indices. Additionally, the EF/PS model fits the data significantly better using a chi-square difference test *χ2*[5] = 20.431, *p* = .001037.

| Model Fit Parameter | EF/PS | COG |
| --- | --- | --- |
| **Akaike Information Criterion (AIC)** | **5016.41** | **5026.84** |
| **Bayesian Information Criterion (BIC)** | **5209.31** | **5203.67** |
| **Root Mean Square Error** | **0.047** | **0.055** |
| **Comparative Fit Index (CFI)** | **0.982** | **0.974** |
| **Tucker-Lewis Index (TLI)** | **0.974** | **0.965** |
| **Standardized Root Mean Square Residual** | **0.03** | **0.036** |
| **Chi-Square Test of Model Fit** | **116.999** | **137.431** |

